# Estimates of introgression as a function of pairwise distances

**DOI:** 10.1101/154377

**Authors:** Bastian Pfeifer, Durrell D Kapan

## Abstract

**Background:** Research over the last 10 years highlights the increasing importance of hybridization between species as a major force structuring the evolution of genomes and potentially providing raw material for adaptation by natural and/or sexual selection. Fueled by research in a few model systems where phenotypic hybrids are easily identified, research into hybridization and introgression (the flow of genes between species) has exploded with the advent of whole-genome sequencing and emerging methods to detect the signature of hybridization at the whole-genome or chromosome level. Amongst these are a general class of methods that utilize patterns of single-nucleotide polymorphisms (SNPs) across a tree as markers of hybridization. These methods have been applied to a variety of genomic systems ranging from butterflies to Neanderthal’s to detect introgression, however, when employed at a fine genomic scale these methods do not perform well to quantify introgression in small sample windows.

**Results:** We introduce a novel method to detect introgression by combining two widely used statistics: pairwise nucleotide diversity *d_xy_* and Patterson’s *D*. The resulting statistic, the *Basic distance fraction* (*Bd_f_*), accounts for genetic distance across possible topologies and is designed to simultaneously detect and quantify introgression. We also relate our new method to the recently published *f_d_* and incorporate these statistics into the powerful genomics R-package PopGenome, freely available on GitHub (*pievos101/PopGenome*). The supplemental material contains a wide range of simulation studies and a detailed manual how to perform the statistics within the PopGenome framework.

**Conclusion:** We present a new distance based statistic *Bd_f_* that avoids the pitfalls of Patterson’s *D* when applied to small genomic regions and accurately quantifies the fraction of introgression (*f*) for a wide range of simulation scenarios.

## Background

Hybridization between species is increasingly recognized as a major evolutionary force. Although long known to occur in plants, evidence is mounting that it regularly occurs in many animal groups [1]. Generally thought to decrease differences between two species by sharing alleles across genomes, hybridization can paradoxically act as a ready source of variation, impacting adaptation [2, 3], aiding in evolutionary rescue [4], promoting range expansion [5], leading to species divergence [6, 7] and ultimately fueling adaptive radiation [8,9]. The advent of whole genome sequencing has prompted the development of a number of methods to detect hybridization across the genome (recently summarized in Payseur and Rieseberg [10])

One class of methods involves quantifying single nucleotide polymorphism (SNP) patterns to detect hybridization between taxa. Here we focus on this class of tests involving four taxa. The most widely used of these, Patterson’s *D*, was first introduced by Green *et al*. [11] and further developed by Durand *et al*. [12]. Patterson’s *D* compares allele patterns of taxa with the Newick tree (((P1,P2),P3),O), to detect introgression between archaic taxon 3 (P3) and in-group taxon 1 (P1) or 2 (P2 or vice-versa). In brief, assuming the outgroup O is fixed for allele A, derived alleles (B) in P3, when shared with either P2 or P1, act as a marker of introgression leading to the following patterns: ABBA or BABA respectively. An excess of either pattern, ABBA or BABA represents a difference from the 50 : 50 ratio expected from incomplete lineage sorting and thus represents a signal that can be used to detect introgression.

Since its introduction, Patterson’s *D* has been used for a wide range of studies to estimate the overall amount of hybrid ancestry by summing the ABBA or BABA pattern excess on a whole genome scale starting with studies of Neanderthals and archaic humans [11, 12]. In the past 7 years, Patterson’s *D* has been increasingly used to localize regions of hybrid ancestry, not only in archaic humans [13] but also in species including butterflies, plants and snakes [14–16].

Currently, Patterson’s *D* is frequently used in sliding window scans of different regions of the genome [17–19]. However, intensive evaluations of the four-taxon ABBA-BABA statistics [20] showed that this approach can lead to many false positives in regions of low recombination and divergence. One of the main reasons is the presence of mainly one of the two alternative topologies as a consequence of a lack of independence of the positions [15], resembling an introgression event, which is exacerbated when analyzing smaller gene-regions. To circumvent this issue, several strategies have been developed. On one side, more sophisticated non-parametric methods have been used to reduce the number of false positives (e.g., Patterson *et al*. [21]). On the other side, new statistics have been developed to better estimate the proportion introgression. Martin *et al*. [20] recently proposed the *f_d_* estimate based on the *f* estimates originally developed by Green *et al*. [11] which measure the proportion of unidirectional introgression from P3 to P2. Specifically, *f_d_* assumes that maximal introgression will lead to equally distributed derived allele frequencies in the donor and the recipient population and therefore utilizes the higher derived allele frequency at each variant site. This strategy aims to model a mixed population maximally affected by introgression. However, this approach has two major shortcomings: First, it is designed to sequentially consider introgression between the archaic population P3 and only one ingroup taxa (P1 or P2). Second, the accuracy of measuring the fraction of introgression strongly depends on the time of gene-flow.

Here we combine the approaches of the four-taxon tests with genetic distance to derive a statistic, the *basic distance fraction* (*Bd_f_*), that estimates the proportion of introgression on a four-taxon tree which strictly ranges from −1 to 1, has symmetric solutions, can be applied to small genomic regions, and is less sensitive to variation in the time of gene-flow than *f_d_*.

## Methods

To derive *Bd_f_* we took a two-fold approach. First, we reformulated Patterson’s D, and *f_d_* in terms of genetic distances based on the hypothesis that past or recent hybridization will leave a signature of reduced *d_xy_* between taxa [18,22]. Second, we account for non-introgressed histories by incorporating distances from species tree patterns into the denominator.

First, following convention, A and B denote ancestral and derived alleles respectively. Derived allele frequencies of the four taxa are *p*_1_*_k_* … *p*_4_*_k_* at variant site *k*. Second, *d_xyk_* is the average pairwise nucleotide diversity between population *x* and *y* at variant site *k*. Each genetic distance can be expressed as a sum of patterns in terms of ancestral and derived alleles allowing the terms ABBA and BABA to be rewritten in terms of genetic distances.

### Patterson’s *D* Statistic as a Function of Pairwise Distances

Here we derive the Patterson’s *D* statistic as a function of pairwise genetic distance between taxon *x* and taxon *y* (*d_xy_*). Following [23] the genetic distance *d_xy_* is defined as

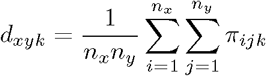
at a given variant site *k*, where *n_x_* is the number of individuals in population *xn_y_* is the number of individuals in population *y*. Then at site *k*, *π_ij_* = 1 ∨ 0 is the boolean value indicating that the individual *i* of population *x* and the individual *j* of population *y* contain the same variant (0) or not (1). The genetic distances *d_xy_* in terms of derived allele frequencies (*p*) are as follows:

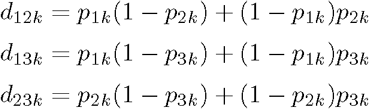

Following [12, 21] instead of pattern counts, allele frequencies can be used as an unbiased estimator. According to that we define *A* as the ancestral allele frequency (1 − *p*) and *B* as the derived allele frequency (*p*) allowing the terms

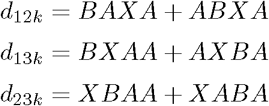
at site *k*. Here X is *A*+*B* = 1 and the position of the letter indicates the population order. The terms ABBA and BABA can then be expressed in terms of distances. If:

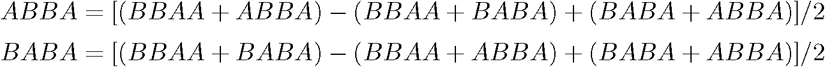
they can be expressed as:

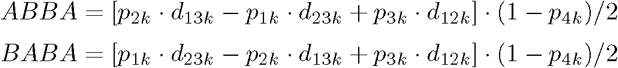

This leads to the following distance based Patterson’s *D* equation for a region containing L variant positions:

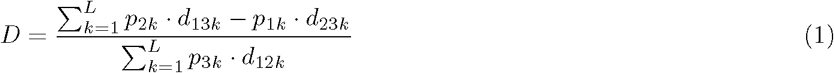
where *d_xyk_* is the average pairwise nucleotide diversity between population *x* and *y* at variant position *k*; and *p_xk_* the derived allele frequency in population *x*. In the context of distances *p*_2_*_k_* · *d*_13_*_k_* may be seen as the contribution of the variation contained between the lineages 1 to 3 (*d*_13_*_k_*) to population 2.

Visualized by equation (1) the Patterson’s *D* denominator (ABBA + BABA) simplifies to an expression of the derived allele frequency of the archaic population P3 times the average pairwise nucleotide diversity (*d_xy_*) between population P1 and P2. This interpretation highlights the original difficulty that Patterson’s *D* has handling regions of low diversity since the denominator will be systematically reduced in these areas due to the *d*_12_*_k_* variable; increasing the overall *D* value. This effect intensifies when at the same time the divergence to the donor population P3 is high. Martin *et al*. [20] proposed *f_d_* which corrects for this by considering the higher derived allele frequency (P2 or P3) at each given variant position; systematically increasing the denominator.

#### Martin’s *f_d_* Estimator

We can apply the same distance logic to rewrite the *f_d_* statistic. Following the example above for *D* we start with the definition of *f_hom_* [11].

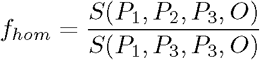
where

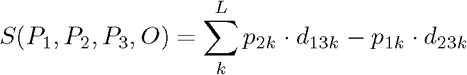

Substituting *P*_2_ with *P*_3_,

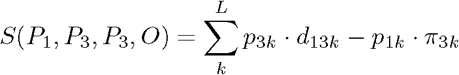
where *π*_3_*_k_* is the average pairwise nucleotide diversity within population P3 at site *k*. *p*_3_*_k_* · *d*_13_*_k_* may be interpreted as the contribution of population 3 to the variation contained between the lineages 1 to 3 (subtracting the contribution of population 1 contained in population 3). Here it is assumed that introgression goes from P3 to P2. Following Martin *et al*. [20] *f_d_* is defined as 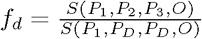 where *P_D_* is the population (2 or 3) with the highest frequency at each variant position. Here the denominator is:

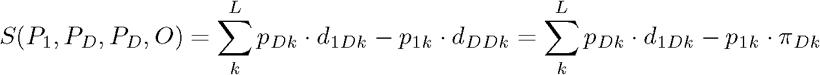

Leading to the statistic:

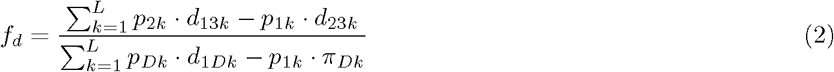
where in the denominator, *π_Dk_* is the average nucleotide diversity within population *P_D_*, which is the population with the higher derived allele frequency in population *P*_2_ or *P*_3_ for each variant site *k*. The difference between the *f_d_* statistic versus *f_hom_* is that there is no assumption in the former about the direction of introgression.

These distance based interpretations suggest there exists a family of related distance estimators for the proportion of introgression. Here we propose a very simple version, we call *Bd_f_*, that makes direct use of the distance based numerator of the Patterson’s *D* statistic and relates the differences of distances to the total distance considered (fig. 1) by incorporating the BBAA species tree pattern into the denominator. The species tree pattern BBAA contributes to increased divergence between (P1,P2) and P3 in the absence of introgression. As a consequence within our *Bd_f_* framework, we explicitly include the divergence to P3 on the four-taxon tree.

**Figure 1.**
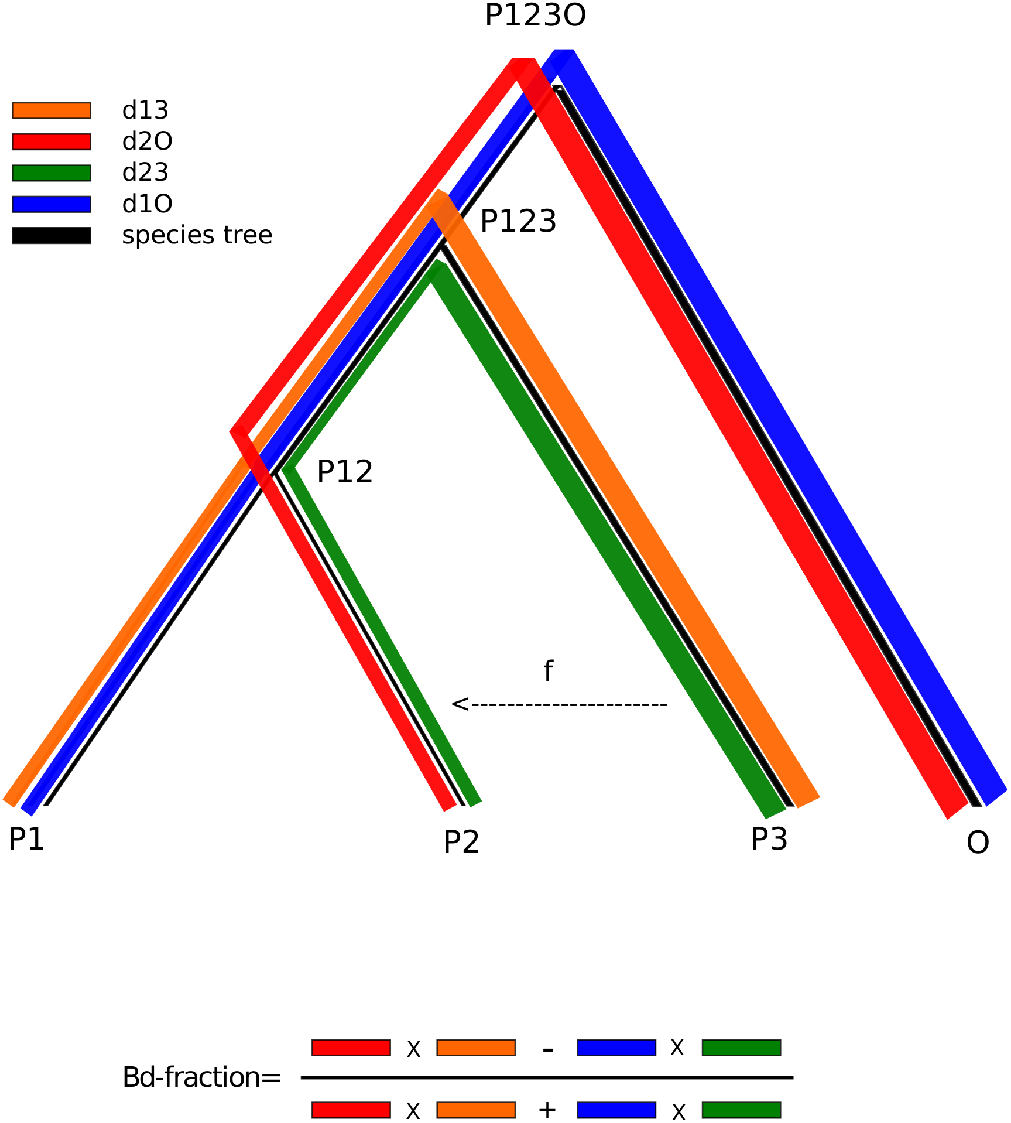
A graphical interpretation of the *Bd_f_* model. A four-taxon tree illustrating the distance based *Bd_f_* model in terms of connecetd path lengths differences. Here *f* is the fraction of introgression from P3 to P2; reducing the distance between P2 and P3 and at the same time increasing the derived allele frequency in P2. *Bd_f_* approximates the measure *f* by relating the differences of the connected path lengths (red, orange) and (blue, green) to the overall sum of connected path lengths. When only sites are considered where the outgroup is mono-allelic the distance to the outgroup (d1O and d2O) simplifies to the derived allele frequencies *p* in population P1 and P2 (blue and red subtree).

#### The *Bd_f_* Estimator

In distance terms we may interpret the ABBA and BABA patterns as polarized shared distances (shared distance between two taxa caused by the derived alleles) on a 4-taxon tree. ABBA for example can be interpreted as the polarized shared distance between (P2,P3) and P1, where BABA is the polarized shared distance between (P1,P3) and P2. Thus, ABBA is a signal of shared increased distance to P1 and BABA is a signal of shared increased distance to P2. However, in order to relate those distances to the distances which are not a signal of introgression, the BBAA pattern must to be taken into account, because the species tree captures the third way in which exactly two populations can share derived alleles. According to the interpretations given above, the BBAA species tree pattern can be seen as the polarized shared distances of (P1,P2) to P3. We incorporate this pattern to refine two classes given the system described above:

- **Class 1:** The contribution of derived alleles in P2 to distance (ABBA+BBAA).
- **Class 2:** The contribution of derived alleles in P1 to distance (BABA+BBAA).

The union of both classes includes all possible patterns producing distances on a 4-taxon tree by shared derived alleles (connected branches in fig. 1). Thus, the denominator of the *Bd_f_* can be written as:

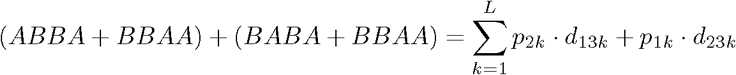

For a given region including *L* variant sites.

A decreased BBAA polarized shared distance and an increased polarized shared distance ABBA is a signal of *P*3 ↔ *P*2 introgression. When at the same time the BABA signal reduces we have a maximal support for the ABBA signal. The *Bd_f_* statistic we propose here has the following form:

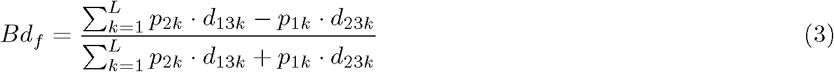

In distance terms, *Bd_f_* may be interpreted as the difference of the distances from P1 and P2 to the archaic population P3 that is caused by introgression. The transformation of the denominator back into the basic Patterson’s *D* statistic form suggests adding the given species tree BBAA pattern to the ABBA and BABA class respectively; which can be reasonably assumed to be the most likely pattern in the absence of introgression for a given species tree (((P1,P2),P3),O). With these patterns in hand it becomes possible to distinguish between signals of introgression and non-introgression. It should be noticed, however, that the *Bd_f_* equation still produces some extreme false positives when e.g the derived allele frequency *p*_1_ or *p*_2_ is zero (often true when block-size is small). Thus, we encourage the user to apply *Laplace smoothing* in genomic scan applications. In this case the derived allele frequency *p* is simply replaced by 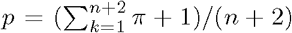 for population 1 & 2 and *d_xy_* is updated accordingly. The parameter *π* is a boolean variable and equals to 1 when a derived allele is present. We have implemented *Laplace smoothing* for *Bd_f_* as a feature in PopGenome.

#### Simulation study

To evaluate the performance of the *Bd_f_* we used a simulation set-up following Martin *et al*. [20]. The Hudson’s ms program [24] was used to generate the topologies with different levels of introgression and the seq-gen program [25] to generate the sequence alignments upon which to compare the performance of the three main statistics discussed in this paper, Patterson’s *D* (*D*), *f_d_* and *Bd_f_* while varying the distance to ancestral populations, time of gene flow, recombination, ancestral population sizes and the effect of low variability. These simulations had the following settings in common: for each fraction of introgression [0, 0.1, …, 0.9, 1], we simulated 100 loci using 5kb windows to calculate three statistics: adjusted *R*^2^ ‘goodness of fit’, The euclidean distance (sum of squared distances) of the mean values to the real fraction of introgression, also called the ‘sum of squares due to lack of fit’ (SSLF) and the ‘pure sum of squares error’ (SSPE). The accuracy of the statistics is shown in fig. 2 and in the supplementary material (tables S1.1-S1.4) for a wide range of simulation parameters.

**Figure 2.**
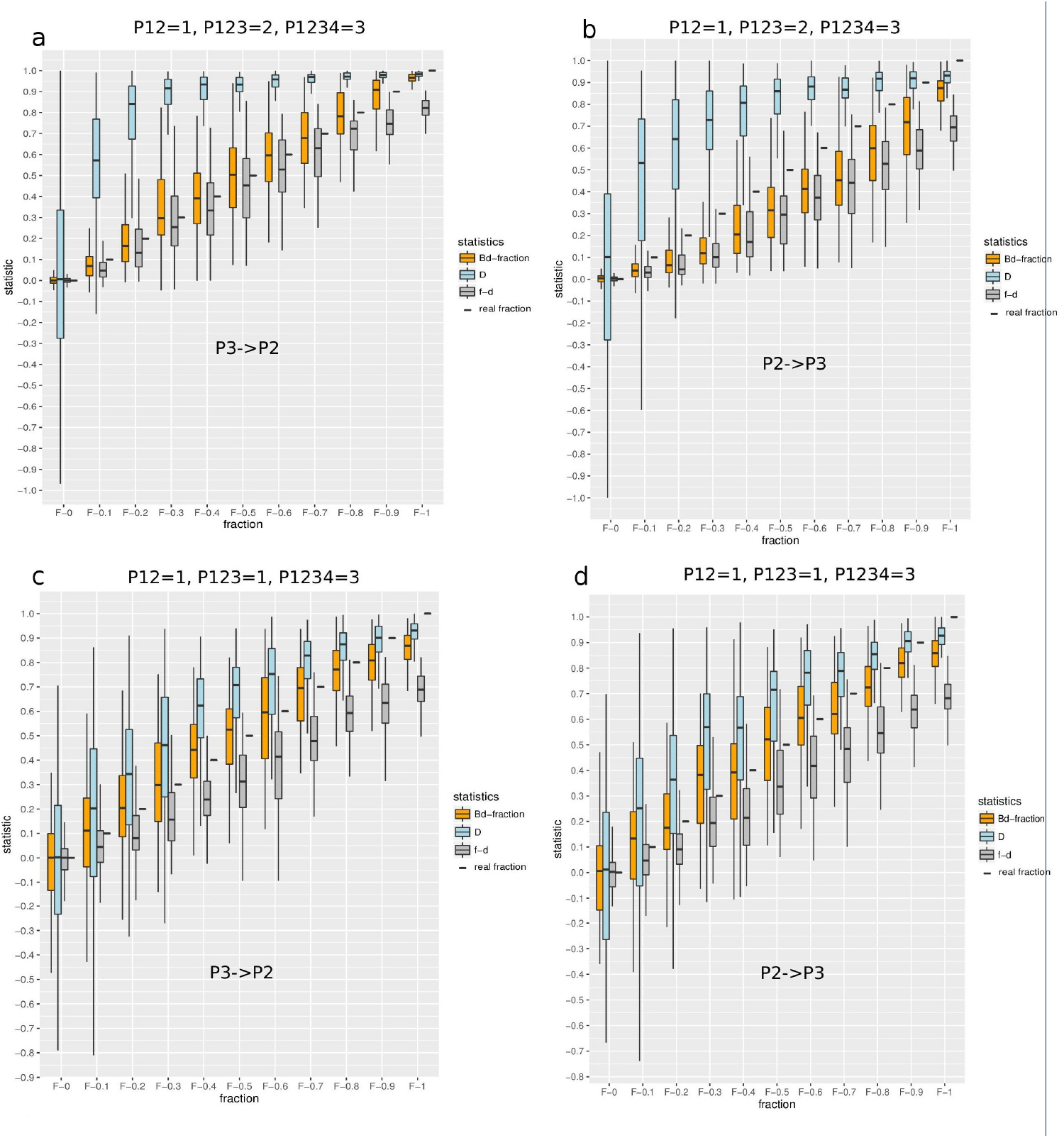
Accuracy of statistics to measure the fraction of introgression. The comparison of simulated data with a known fraction of introgression using ms versus the statistics (y-axis). We simulated 100 loci for every fraction of introgression [0, 0.1, … 0.9, 1] and plotted the distribution of the corresponding statistic outcomes. A window size of 5kb and a recombination rate of r=0.01 was used. The background histories (coalescent events) are **a:** P12=1*N_e_*, P123=2*N_e_*, P1234=3*N_e_* generations ago. **b:** P12=1*N_e_*, P123=2*N_e_*, P1234=3*N_e_* generations ago. **c:** P12=1*N_e_*, P123=1*N_e_*, P1234=3*N_e_* generations ago. **d:** P12=1*N_e_*, P123=1*N_e_*, P1234=3*N_e_* generations ago. Introgression directions are *P*3 → *P*2 (A,C) and *P*2 → *P*3 (B,D). Colors: *f_d_* (grey), *Bd_f_* (orange) Patterson’s *D* (light blue) and the real fraction of introgression (black boxes).

All of these analyses were done in the R-package PopGenome [26], that efficiently calculates *Bd_f_* (and other statistics including *f_d_*, *RNDmin* [27], and the original Patterson’s *D*) from the scale of individual loci to entire genomes.

## Results

We performed extensive simulations varying the distance to ancestral populations, time of gene flow, recombination, ancestral population sizes and mutation rates. We found that *Bd_f_* outperforms or is essentially equivalent to the *f_d_* estimate to measure the real fraction of introgression for most of the studied ranges of simulation cases. Overall, because it captures natural variation in the denominator, *Bd_f_* has slightly higher variances compared to *f_d_* while the mean values are often the least biased as shown by the sum of squares due to lack of fit, yet it provides the best (or nearly equivalent) estimates to *f_d_* as judged by the goodness of fit in almost all cases (supplementary information, section S1).

### The effect of background history

Simulations under a variety of coalescent times show that *Bd_f_* is the most accurate approximation of the real fraction of introgression, including under the different coalescent events simulated for both directions of introgression (fig. 2, table 1). Following behind *Bd_f_* is *f_d_*, which is more affected by differences in coalescent times. In this comparison, Patterson’s *D* consistently overestimates the fraction of introgression (fig. 2, table 1). This known effect [20] is greatest in the most common case where the coalescent times differ between ingroup taxa (P1,P2) and the archaic taxon P3. This effect is also slightly impacted by the direction of introgression (fig. 2, table 1). However, for the more unrealistic case where the ingroup taxa (P1,P2) and the archaic taxon P3 are evolutionary very close it should be noticed that *Bd_f_* essentially differs from the *f_d_* estimate. In this specific case the ‘pure sum of squares error’ (SSPE) of *Bd_f_* increases leading to a lower ‘goodness of fit’ value compared to *f_d_*, while the ‘sum of squares due to lack of fit’ (SSLF) are still notably low signifying a very precise mean estimate of the real fraction of introgression. From Figure 2 we see that the *Bd_f_* related SSPE values are high only if the signal of introgression is very low. So, we expect *Bd_f_* to quantify stronger signals of introgression more precisely.

**Table 1.** The effect of the distance to ancestral population. This table refers to Figure 2 and displays some supporting values.

| Direction of gene-flow | Distance to ancestral | $D$ | $f_d$ | $Bd_f$ | |
| --- | --- | --- | --- | --- | --- |
| $P3 \rightarrow P2$ | 1-1-3 | 0.58 | 0.77 | 0.70 | a |
|  |  | 0.12 | 0.40 | 0.04 | b |
|  |  | 0.60 | 0.17 | 0.35 | c |
| $P3 \rightarrow P2$ | 1-2-3 | 0.39 | 0.80 | 0.81 | a |
|  |  | 1.41 | 0.09 | 0.00 | b |
|  |  | 0.48 | 0.19 | 0.25 | c |
| $P2 \rightarrow P3$ | 1-1-3 | 0.57 | 0.76 | 0.70 | a |
|  |  | 0.12 | 0.42 | 0.05 | b |
|  |  | 0.59 | 0.17 | 0.33 | c |
| $P2 \rightarrow P3$ | 1-2-3 | 0.40 | 0.78 | 0.77 | a |
|  |  | 0.70 | 0.54 | 0.30 | b |
|  |  | 0.48 | 0.19 | 0.19 | c |

<sup>a</sup> the adjusted $R^2$ 'goodness of fit' (*higher is better*).
<sup>b</sup> SSLF 'sum of squares due to lack of fit' divided by the sample size $n=100$ (*lower is better*).
<sup>c</sup> SSPE 'pure sum of squares error' (*lower is better*).

### The effect of the time of gene-flow

One advantage of *Bd_f_* compared to the other methods studied in this paper is that it is rarely affected by the time of gene-flow (fig. 3). This is due to the fact that, unlike *f_d_*, *Bd_f_* does not relate the signal of introgression to its maximum calculated from the present. When gene flow occurs in the distant past the denominator of *f_d_* estimates increases leading to an underestimation of the fraction of introgression. The ‘goodness of fit’ of *Bd_f_* is consistently higher than *f_d_* (fig 3A), but more importantly, at the same time the SSLF values are almost unaffected by the time of gene-flow (fig. 3B). Notably, the direction of gene-flow has an effect that synergizes with the time that it occurred, with introgression between *P*2 → *P*3 in the distant past overall showing lower values of the statistics.

**Figure 3.**
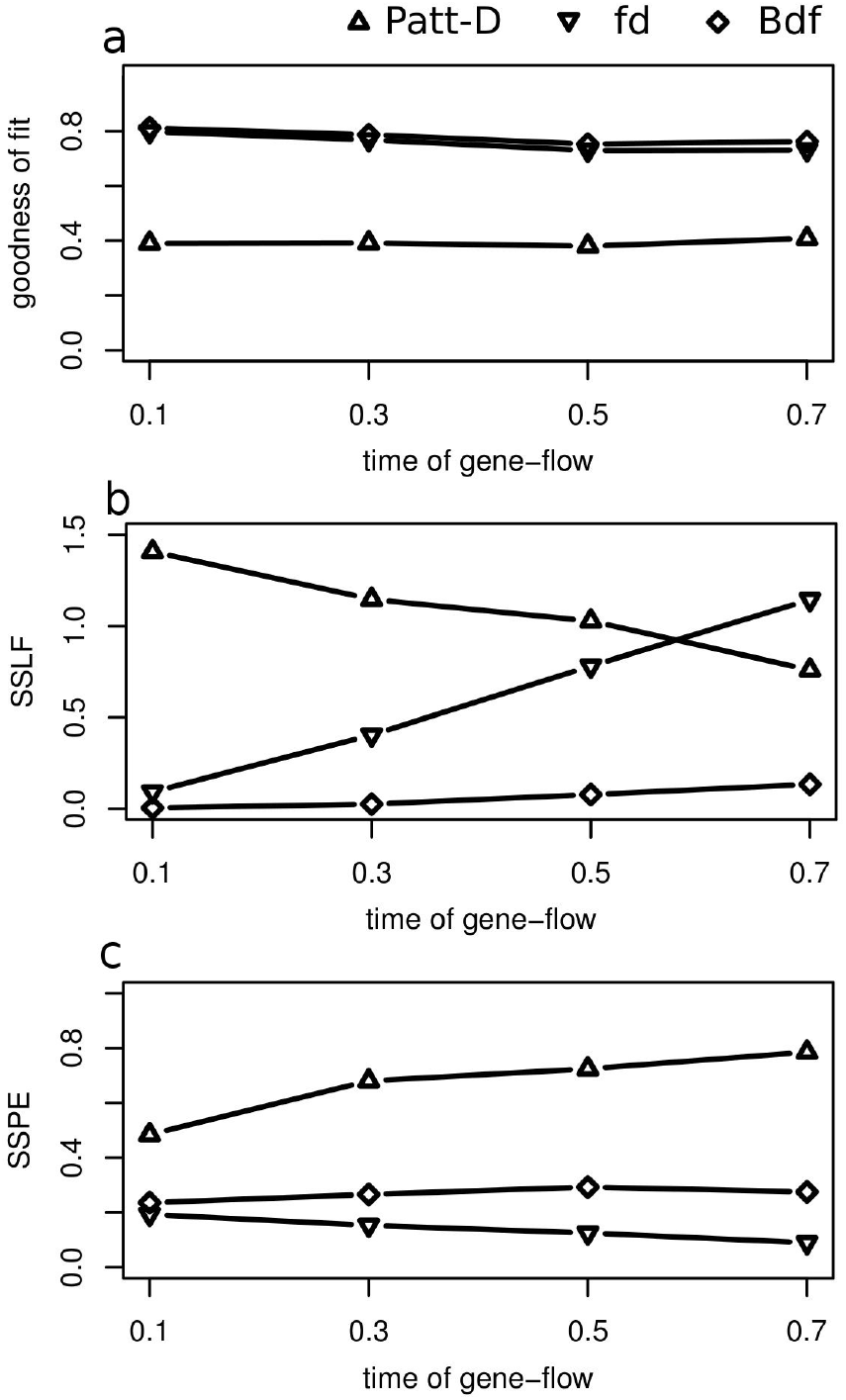
The effect of time of gene-flow. For *P*3 → *P*2 introgression we varied the time of gene-flow (0.1, 0.3, 0.5, 0.7 *N_e_*) and calculated for each statistic (*D*, *f_d_* and *Bd_f_*) **a:** the adjusted *R*^2^ ‘goodness of fit’. **b:** SSLF ‘sum of squares due to lack of fit’ divided by the sample size n=100. **c:** SSPE ‘pure sum of squares error’. Scaled recombination rate is *N_e_r*=50 (*r* = 0.01). The background history is: P12=1*N_e_*, P123=2*N_e_* and P1234=3*N_e_* generations ago. The calls to ms are: *P*3 → *P*2: ms 32 1 -I 4 8 8 8 8 -ej 1 2 1 -ej 2 3 1 -ej 3 4 1 -es Gene-flow 2 Fraction -ej Gene-flow 5 3 -r 50 5000. *P*2 → *P*3: ms 32 1 -I 4 8 8 8 8 -ej 1 2 1 -ej 2 3 1 -ej 3 4 1 -es Gene-flow 3 Fraction -ej Gene-flow 5 2 -r 50 5000

### The effect of recombination

We found that all three methods *Bd_f_*, *f_d_* and Patterson’s *D* become more accurate with increasing recombination rates. This is due to the increase of independent sites of a region analyzed. While *Bd_f_* tends to have higher variances when the recombination rate is low it’s variance is comparable to *f_d_* as soon as the recombination rate increases (supplementary table S1.2).

### On the ability to detect introgression

To further test *Bd_f_*, we evaluated the performance to detect introgression by simulating 10,000 neutral loci and 10 loci subject to introgression, interpreting the results using a receiver operating characteristic curve (ROC) analysis that evaluates the area under the curve (AUC) a measure that summarizes model performance, the ability to distinguish introgression from the neutral case, calculated with the R-package pROC [28]. For this simulation scenario *Bd_f_* and the *f_d_* estimate show nearly the same utility (higher is better) for the fraction of introgression and distance to ancestral population (supplementary information, section S2); but both, in agreement with Martin *et al*. [20], greatly outperform the Patterson’s *D* statistic especially for smaller genomic regions. We also included the recently published *RNDmin* method in this latter analysis; this alternative only gives good results when the signal of introgression is very strong (supplementary information, section S2). In addition, unlike *f_d_*, *Bd_f_* is able to quantify the proportion of admixture symmetrically (*P*3 ↔ *P*2 and *P*3 ↔ *P*1) it simplifies the analysis of real genomic data on a 4-taxon system.

### Application

To test with real data we calculated *Bd_f_* for 50kb consecutive windows on the 3L arm of malaria vectors in the *Anopheles gambiae* species complex (fig. 4) confirming the recently detected region of introgression found in an inversion [17]. In order to detect chromosome-wide outliers we tested the null hypotheses (*Bd_f_* = 0) *outside* of the inversions and *inside* the inversion 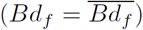. The analyses was done on the basis of 50 kb consecutive windows using a weighted block jackknife to generate Z-values. The corresponding *P* values were corrected by multiple testing using the Benjamin-Hochberg false discovery rate (FDR) method [29].

**Figure 4.**
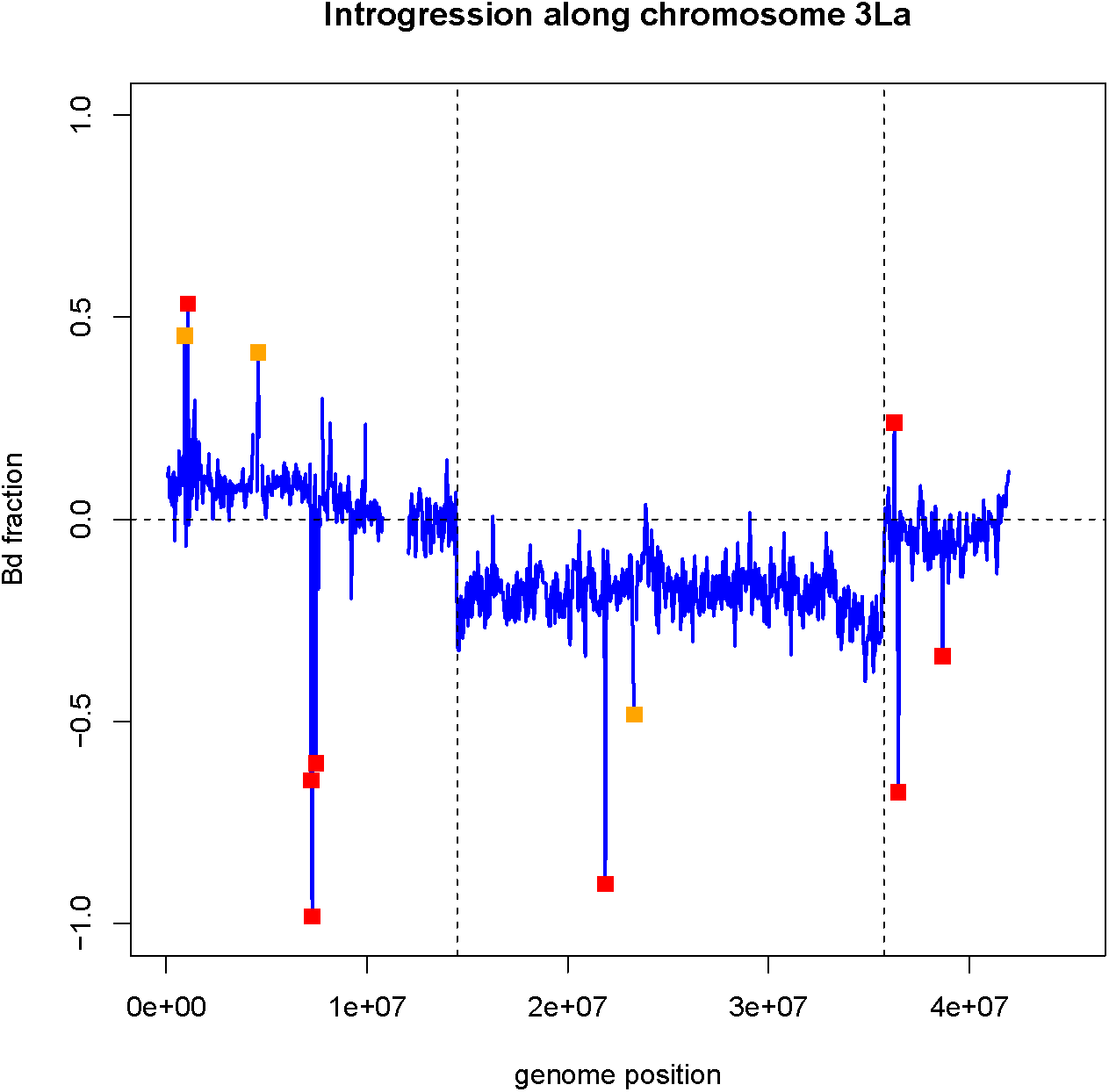
Anopheles gambiae 3La inversion. Confirming introgression on the 3L arm of the malaria vector Anopheles gambiae (Fontaine *et al*. 2015, fig. 4). We used the R-package PopGenome to scan the chromosome with 50kb consecutive windows and plotted the *Bd_f_* values along the chromosome (*Laplace smoothed*). Orange boxes indicate outlier windows below a significance level of 0.05 and red boxes show outlier windows on the basis of a 0.01 significance level. The p-values were corrected for multiple testing by the Benjamin-Hochberg method.

Overall, we found 9 significant outliers outside the inversion and two outliers within the inversion based on a 0.05 significance level (see figure 4). This further reduces to 7 significant outliers outside the inversion and one remaining outlier within the inversion when tested against a 0.01 significance level (see table 2).

**Table 2.**
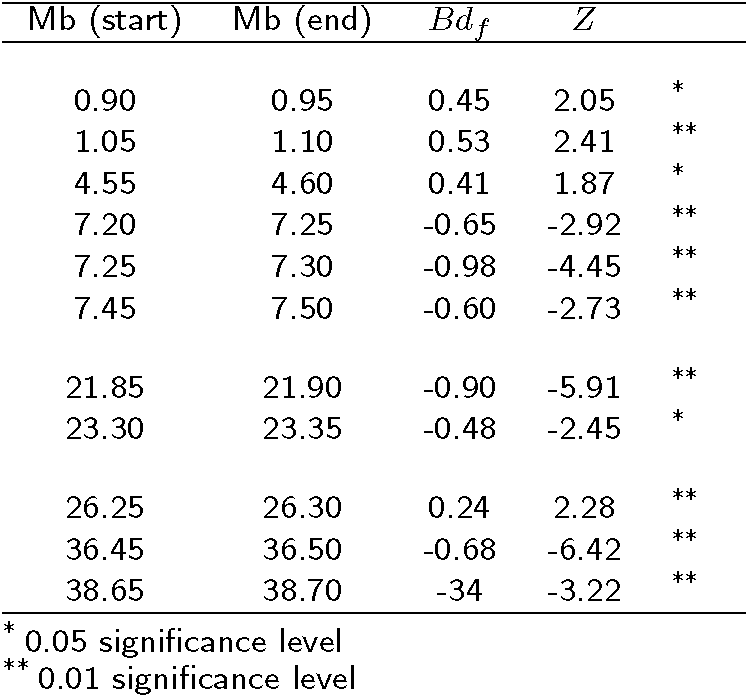
Significant outlier detected on the Anopheles gambiae 3La chromosome

These analyses were all performed within the R package PopGenome [26] and can be easily reproduced with the code given in the supplementary material section S3.

## Discussion

In the last 8 years there has been an explosion of population genomic methods to detect introgression. The Patterson’s *D* method, based on patterns of alleles in a four-taxon comparison, has been widely applied to a variety of problems that differ from those for which it was originally developed. This statistic can be used to assess whether or not introgression is occurring at the whole genome scale, however, Patterson’s *D* is best not applied to smaller genomic regions or gene-scans as noted by Martin *et al*. 2015.

The distance based approach proposed here has the following strengths: First, the distance approach points to a family of statistics that can directly identify changes in genetic distances that are a natural consequence of introgression. Second, distance measured by *d_xy_* allows direct comparisons of quantities that are easily interpreted. Third, a simple member of this family based on these distances, *Bd_f_*, accurately predicts the fraction of introgression over a wide-range of simulation parameters. Furthermore, the *Bd_f_* statistic is symmetric (like Patterson’s D) which makes it easy to implement and interpret. However, *Bd_f_* also outperforms Patterson’s *D* in all cases (the latter shows a strong positive bias) and *Bd_f_* also outperforms or is equivalent to *f_d_* in nearly all cases judged by the goodness of fit and the sum of squares due to lack of fit. Furthermore, unlike *f_d_*, *Bd_f_* does not vary strongly with the time of gene-flow. This latter strength comes from incorporating the genetic distance to taxon 3 (P3) into the denominator, serving to scale *Bd_f_* relative to *d_xy_* values between the three species in the comparisons. Ultimately this makes he statistic *less* subject to extreme false positives due to low SNP diversity (low genetic distances), as evidence by lower values than other statistics in our examples.

There are several areas where further improvements could be made. Although the distance based derivation of all three statistics is sound, and *Bd_f_* is empirically supported by simulation, further mathematical analysis for this general class of distance estimators is desired. Like other statistics under consideration in this paper, *Bd_f_* depends on resolved species tree with a particular configuration of two closely related species, a third species and an outgroup, and therefore it is not directly applicable to other scenarios. In addition, both the *f_d_* and *Bd_f_* perform less accurately when measuring the proportion of admixture when the gene-flow occurs from P2 to P3. On the other hand, our simulations revealed (Figure 5) the asymmetrical affect of gene-flow direction on genetic distance: gene-flow from P3 to P2 does not affect the distance between taxon 1 & 3 (*d*_13_), however, the opposite it true when introgression from P2 to P3 occurs, the distance between taxon 1 & 2 (*d*_12_) is not affected. This suggests comparisons of *d_xy_* within given genomic regions may contain signal to infer the direction of introgression and therefore more accurately measure the proportion of admixture.

**Figure 5.**
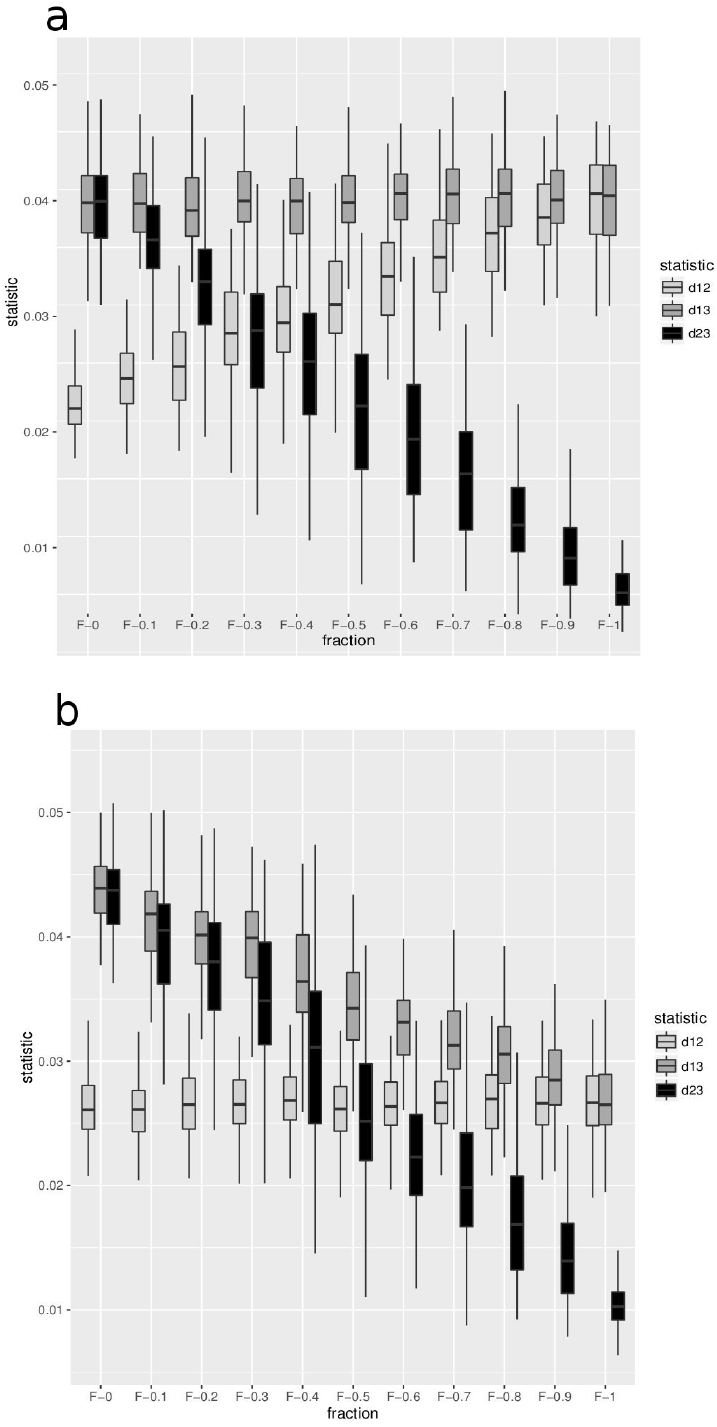
The effect of introgression on pairwise distances. The effect of the fraction of introgression on the average pairwise distance measurements *d*_12_, *d*_13_ and *d*_23_. **a:** The effect is shown for *P*3 → *P*2 introgression. **b:** Shows the effect in case of *P*2 → *P*3 introgression. The background history is: P12=1*N_e_*, P123=2*N_e_* and P1234=3*N_e_* generations ago.

Overall, the distance based interpretation of introgression statistics suggests a general framework for estimation of the fraction of introgression on a known tree and may be extended in a few complementary directions including the use of model based approaches to aid in outlier identification and potentially model selection. The distance based framework introduced here may lead to other further improvements by measuring how genetic distance changes between different taxa as a function of hybridization across different parts of the genome.

## Conclusion

Here we present both a simplified distance based interpretation for Patterson’s *D* and Martin *et al.’s f_d_* and a new distance based statistic *Bd_f_* that avoids the pitfalls of Patterson’s *D* when applied to small genomic regions and is more accurate and less prone to vary with variation in the time of gene flow than *f_d_*. We propose *Bd_f_* as an estimate of introgression which can be used to simultaneously detect and quantify introgression. We implement *Bd_f_* (as well as the other four-taxon statistics, *f_d_*, and the original Patterson’s *D*) in the powerful R-package, PopGenome [26], now updated to easily calculate these statistics for individual loci to entire genomes.

## Acknowledgements

We would like to thank Bettina Harr, Matthew Hansen, Jim Henderson, Karl Lindberg, Paul Staab, Sebastian E. Ramos-Onsins, the Academy genomics discussion group and the IMI journal club for helpful discussions.

## Availability of data and materials

An updated PopGenome package including the methods presented in this paper is available for download from a GitHub repository (https://github.com/pievos101/PopGenome). R-code to reproduce the simulations can be found at https://github.com/pievos101/Introgression-Simulation.

## Author’s contributions

BP and DDK designed the project. BP developed the methods and performed the simulations. BP and DDK wrote the manuscript.

