## Supplementary Materials for "Estimates of introgression as a function of pairwise distances" (from another version of this preprint: version 4, 2018-03-15)

#### 1 On the Accuracy to Measure the Real Fraction of Introgression

Distance of ancestral population: Starting with the following topology (((P1,P2),P3),P4) we simulate varying depths to common ancestors of P1 & P2, and at the root (P1234), where recombination rate is fixed with $N_e r$ =50 ($r$=0.01) for each direction of gene flow.

**Supplementary Table 1.** Distance of ancestral population.

| Direction of gene-flow | Distance to ancestral population (P12-P123-P1234) | *D* | $f_d$ | $Bd_f$ | |
|---|---|---|---|---|---|
| P3→P2 | 0.3-1-3 | 0.4388 | 0.7662 | 0.7739 | a |
| | | 0.7060 | 0.2893 | 0.1884 | b |
| | | 0.5175 | 0.1667 | 0.2210 | c |
| P3→P2 | 0.5-1-3 | 0.4916 | 0.7622 | 0.7837 | a |
| | | 0.4895 | 0.3283 | 0.1278 | b |
| | | 0.5619 | 0.1676 | 0.2180 | c |
| P3→P2 | 0.7-1-3 | 0.5158 | 0.7675 | 0.7698 | a |
| | | 0.2928 | 0.3667 | 0.0733 | b |
| | | 0.6544 | 0.1695 | 0.2487 | c |
| P3→P2 | 1-1-3 | 0.5846 | 0.7733 | 0.7014 | a |
| | | 0.1177 | 0.4003 | 0.0380 | b |
| | | 0.6033 | 0.1667 | 0.3540 | c |
| P3→P2 | 0.5-2-3 | 0.3314 | 0.7901 | 0.8064 | a |
| | | 1.6221 | 0.0814 | 0.0798 | b |
| | | 0.6026 | 0.2005 | 0.2252 | c |
| P3→P2 | 0.5-3-3 | 0.2704 | 0.8048 | 0.8077 | a |
| | | 1.9736 | 0.0299 | 0.0884 | b |
| | | 0.6129 | 0.2087 | 0.2255 | c |

| | | | | | |
|---|---|---|---|---|---|
| P3→P2 | 1-2-3 | 0.3905 | 0.7978 | 0.8115 | a |
| | | 1.4075 | 0.0928 | 0.0048 | b |
| | | 0.4839 | 0.1931 | 0.2361 | c |
| P3→P2 | 1-3-3 | 0.3267 | 0.8142 | 0.8212 | a |
| | | 1.7027 | 0.0343 | 0.0269 | b |
| | | 0.6096 | 0.2039 | 0.2297 | c |
| P3→P2 | 2-2-3 | 0.5858 | 0.8115 | 0.7254 | a |
| | | 0.4688 | 0.1920 | 0.1339 | b |
| | | 0.4950 | 0.1761 | 0.3274 | c |
| P2→P3 | 0.3-1-3 | 0.1593 | 0.4724 | 0.4320 | a |
| | | 0.8882 | 1.8376 | 2.2058 | b |
| | | 0.7641 | 0.1412 | 0.1072 | c |
| P2→P3 | 0.5-1-3 | 0.3660 | 0.6895 | 0.6465 | a |
| | | 0.2165 | 1.0807 | 1.0442 | b |
| | | 0.7532 | 0.1314 | 0.1662 | c |
| P2→P3 | 0.7-1-3 | 0.4925 | 0.7283 | 0.7001 | a |
| | | 0.1415 | 0.6811 | 0.3466 | b |
| | | 0.6317 | 0.1551 | 0.2418 | c |
| P2→P3 | 1-1-3 | 0.5745 | 0.7569 | 0.7040 | a |
| | | 0.1211 | 0.4169 | 0.0473 | b |
| | | 0.5876 | 0.1742 | 0.3340 | c |
| P2→P3 | 0.5-2-3 | 0.2525 | 0.6755 | 0.6183 | a |
| | | 0.3321 | 1.2255 | 1.4610 | b |
| | | 0.8886 | 0.1285 | 0.1333 | c |
| P2→P3 | 0.5-3-3 | 0.2361 | 0.6709 | 0.5852 | a |
| | | 0.3760 | 1.3214 | 1.6903 | b |
| | | 0.9414 | 0.1249 | 0.1160 | c |
| P2→P3 | 1-2-3 | 0.3952 | 0.7778 | 0.7691 | a |
| | | 0.6901 | 0.5376 | 0.3026 | b |
| | | 0.4839 | 0.1931 | 0.1895 | c |
| P2→P3 | 1-3-3 | 0.3767 | 0.7589 | 0.7226 | a |
| | | 0.8946 | 0.6210 | 0.6119 | b |
| | | 0.6854 | 0.1570 | 0.1924 | c |
| P2→P3 | 2-2-3 | 0.5628 | 0.8020 | 0.7188 | a |
| | | 0.4822 | 0.1948 | 0.1419 | b |
| | | 0.5399 | 0.1839 | 0.3309 | c |

For each direction of gene flow and distance to ancestral populations (see above) we calculated for each statistic (*D*, $f_d$ and $Bd_f$) [a] the adjusted $R^2$ 'goodness of fit'. [b] *SSLF* 'sum of squares due to lack of fit' divided by the sample size n=100. [c] SSPE 'pure sum of squares error'. The time of gene-flow was a constant at 0.1$N_e$, the scaled recombination rate is $N_e r$=50 ($r$=0.01), and the calls to ms are as follows:

P3→P2: ms 32 1 -I 4 8 8 8 8 -ej P12 2 1 -ej P123 3 1 -ej P1234 4 1 -es 0.1 2 Fraction -ej 0.1 5 3 -r

50 5000

P2→P3: ms 32 1 -I 4 8 8 8 8 -ej P12 2 1 -ej P123 3 1 -ej P1234 4 1 -es 0.1 3 Fraction -ej 0.1 5 2 -r 50 5000

**Supplementary Table 2.** Effect of the time of gene-flow.

| Direction of gene-flow | Time of Gene-flow | *D* | $f_d$ | $Bd_f$ | |
|---|---|---|---|---|---|
| P3→P2 | 0.1 | 0.3905 | 0.7978 | 0.8115 | a |
| | | 1.4075 | 0.0928 | 0.0048 | b |
| | | 0.4839 | 0.1931 | 0.2361 | c |
| P3→P2 | 0.3 | 0.3918 | 0.7681 | 0.7870 | a |
| | | 1.1462 | 0.4039 | 0.0255 | b |
| | | 0.6805 | 0.1530 | 0.2659 | c |
| P3→P2 | 0.5 | 0.3805 | 0.7291 | 0.7525 | a |
| | | 1.0277 | 0.7815 | 0.0782 | b |
| | | 0.7240 | 0.1250 | 0.2924 | c |
| P3→P2 | 0.7 | 0.4084 | 0.7308 | 0.7620 | a |
| | | 0.7601 | 1.1447 | 0.1341 | b |
| | | 0.7850 | 0.0895 | 0.2751 | c |
| P2→P3 | 0.1 | 0.3952 | 0.7778 | 0.7691 | a |
| | | 0.6901 | 0.5376 | 0.3026 | b |
| | | 0.4839 | 0.1931 | 0.1895 | c |
| P2→P3 | 0.3 | 0.3702 | 0.6938 | 0.7003 | a |
| | | 0.4257 | 1.2463 | 0.5155 | b |
| | | 0.8134 | 0.1078 | 0.2379 | c |
| P2→P3 | 0.5 | 0.3069 | 0.5779 | 0.5936 | a |
| | | 0.2800 | 2.0306 | 0.9803 | b |
| | | 0.9632 | 0.0690 | 0.2356 | c |
| P2→P3 | 0.7 | 0.1639 | 0.4283 | 0.4617 | a |
| | | 0.4376 | 2.7323 | 1.7089 | b |
| | | 1.3510 | 0.0432 | 0.1957 | c |

For each direction of introgression we varied the time of gene-flow (0.1, 0.3, 0.5, 0.7 $N_e$) and calculated for each statistic (*D*, $f_d$ and $Bd_f$) [a] the adjusted $R^2$ 'goodness of fit'. [b] SSLF 'sum of squares due to lack of fit' divided by the sample size n=100. [c] SSPE 'pure sum of squares error'. Scaled recombination rate is $N_e r=50$ (r=0.01). The background history is: P12=1$N_e$, P123=2$N_e$ and P1234=3$N_e$ generations ago. The calls to ms are:

*P3→P2: ms 32 1 -I 4 8 8 8 8 -ej 1 2 1 -ej 2 3 1 -ej 3 4 1 -es **Gene-flow** 2 **Fraction** -ej **Gene-flow** 5 3 -r 50 5000*

Recombination: To test the impact of recombination on these statistics we varied the recombination rates from ($r$ = 0 - .08). With increasing recombination rates the accuracy to measure the real fraction of introgression increases for $f_d$ and $Bd_f$ while the Patterson's $D$ is rarely affected by varying this parameter.

**Supplementary Table 3.** Recombination.

| Direction of gene-flow | Recombination rate | *D* | $f_d$ | $Bd_f$ | |
|---|---|---|---|---|---|
| P3→P2 | 0/5000 | 0.3307 | 0.6335 | 0.6062 | a |
| | | 0.7697 | 0.1066 | 0.0139 | b |
| | | 1.4885 | 0.4485 | 0.6535 | c |
| P3→P2 | 50/5000 | 0.3905 | 0.7978 | 0.8115 | a |
| | | 1.4075 | 0.0928 | 0.0048 | b |
| | | 0.4839 | 0.1931 | 0.2361 | c |
| P3→P2 | 100/5000 | 0.4072 | 0.8676 | 0.8758 | a |
| | | 1.4905 | 0.1114 | 0.0080 | b |
| | | 0.3725 | 0.1155 | 0.1467 | c |
| P3→P2 | 200/5000 | 0.4046 | 0.8986 | 0.9020 | a |
| | | 1.5117 | 0.1163 | 0.0121 | b |
| | | 0.3639 | 0.0864 | 0.1096 | c |
| P3→P2 | 300/5000 | 0.4069 | 0.9249 | 0.9257 | a |
| | | 1.5641 | 0.1209 | 0.0150 | b |
| | | 0.3191 | 0.0625 | 0.0820 | c |
| P3→P2 | 400/5000 | 0.4041 | 0.9383 | 0.9360 | a |
| | | 1.5778 | 0.1223 | 0.0169 | b |
| | | 0.3186 | 0.0510 | 0.0691 | c |

For recombination rates varying from (0 to .08) we calculated for each statistic ($D$, $f_d$ and $Bd_f$) [a] the adjusted $R^2$ 'goodness of fit'. [b] *SSLF* 'sum of squares due to lack of fit' divided by the sample size n=100 . [c] SSPE 'pure sum of squares error'. The time of gene-flow was a constant at $0.1N_e$, background history is P12=$1N_e$, P123=$2N_e$ and P1234=$3N_e$, and the calls to ms are as follows:

*P3→P2: ms 32 1 -I 4 8 8 8 8 -ej 1 2 1 -ej 2 3 1 -ej 3 4 1 -es 0.1 2* ***Fraction*** *-ej 0.1 5 3 -r* ***Ne*r*** *5000*

*only one direction of gene-flow shown.*

Ancestral population sizes: We varied ancestral population sizes at the nodes P12 and P123 and simulated the impact on the ($D$, $f_d$ and $Bd_f$) statistics (see supplementary table 4 below).

**Supplementary Table 4.** Ancestral population sizes.

| Direction of gene-flow | Ancestral population size P12-P123 | $D$ | $f_d$ | $Bd_f$ | |
|---|---|---|---|---|---|
| P3→P2 | 1-2 | 0.4133 | 0.8275 | 0.8342 | a |
| | | 1.1476 | 0.0668 | 0.0304 | b |
| | | 0.6070 | 0.1809 | 0.2065 | c |
| P3→P2 | 1-10 | 0.4606 | 0.8031 | 0.8062 | a |
| | | 1.0477 | 0.0084 | 0.1009 | b |
| | | 0.4794 | 0.2411 | 0.2165 | c |
| P3→P2 | 2-10 | 0.5955 | 0.7950 | 0.8022 | a |
| | | 0.5244 | 0.0095 | 0.0474 | b |
| | | 0.4276 | 0.2555 | 0.2420 | c |
| P3→P2 | 2-1 | 0.4857 | 0.8177 | 0.8237 | a |
| | | 0.9736 | 0.1292 | 0.0036 | b |
| | | 0.4743 | 0.1743 | 0.2197 | c |
| P3→P2 | 10-1 | 0.5790 | 0.8000 | 0.7677 | a |
| | | 0.6137 | 0.1867 | 0.0649 | b |
| | | 0.4147 | 0.1841 | 0.2693 | c |
| P3→P2 | 10-2 | 0.6208 | 0.7957 | 0.7778 | a |
| | | 0.4066 | 0.1200 | 0.0147 | b |
| | | 0.4288 | 0.2109 | 0.2804 | c |

For different ancestral population sizes (multiples of 1, 2 and 10$N_e$) at nodes P12 and P123 we calculated for each statistic ($D$, $f_d$ and $Bd_f$) and present [a] the adjusted $R^2$ 'goodness of fit'. [b] *SSLF* 'sum of squares due to lack of fit' divided by the sample size n=100. [c] SSPE 'pure sum of squares error'. The time of gene-flow is 0.1$N_e$, scaled recombination rate is $N_e r$=50 ($r$=0.01), and background history is: P12=1$N_e$, P123=2$N_e$ and P1234=3$N_e$, and the calls to *ms* are:

*P3→P2: ms 32 1 -I 4 8 8 8 8 -ej 1 2 1 -en 1.01 1* ***P12*** *-ej 2 3 1 -en 2.01 1* ***P123*** *-ej 3 4 1 -es 0.1 2* ***Fraction*** *-ej 0.1 5 3 -r 50 5000*

The effect of low variability: We varied the nucleotide diversity θ to test the effect of low variability on the statistics *D*, $f_d$ and $Bd_f$.

**Supplementary Table 5.** The effect of low variability.

| Direction of gene-flow | Variability theta (θ) | *D* | $f_d$ | $Bd_f$ | |
|---|---|---|---|---|---|
| P3→P2 | 3/5000 | 0.3960 | 0.8153 | 0.8197 | a |
| | | 1.3084 | 0.1108 | 0.0130 | b |
| | | 0.5743 | 0.1772 | 0.2252 | c |
| P3→P2 | 5/5000 | 0.4015 | 0.8083 | 0.8205 | a |
| | | 1.3239 | 0.1218 | 0.0192 | b |
| | | 0.5362 | 0.1832 | 0.2206 | c |
| P3→P2 | 25/5000 | 0.4035 | 0.7991 | 0.8102 | a |
| | | 1.2020 | 0.1310 | 0.0229 | b |
| | | 0.6193 | 0.1923 | 0.2379 | c |
| P3→P2 | 50/5000 | 0.4092 | 0.8068 | 0.8217 | a |
| | | 1.3574 | 0.1009 | 0.0064 | b |
| | | 0.4953 | 0.1866 | 0.2277 | c |

For unidirectional gene-flow P3→P2, we varied θ from 3/5000 - 50/5000 and calculated for each statistic (*D*, $f_d$ and $Bd_f$) and present [a] the adjusted $R^2$ 'goodness of fit'. [b] *SSLF* 'sum of squares due to lack of fit' divided by the sample size n=100. [c] SSPE 'pure sum of squares error'. The time of gene-flow was a constant at $0.1N_e$, the scaled recombination rate is $N_e*r$=50 (*r*=0.01) and background history is P12=$1N_e$, P123=$2N_e$ and P1234=$3N_e$. The calls to *ms* are:

*P3→P2: ms 32 1 -I 4 8 8 8 8 -ej 1 2 1 -ej 2 3 1 -ej 3 4 1 -es 0.1 2* ***Fraction*** *-ej 0.1 5 3 -r 50 5000 -t* ***variability***

### 2 Detecting Introgression from Whole Genome Data

To test the performance of the various statistics (*D*, $f_d$, *RNDmin*, $Bd_f$) to distinguish neutral models from models with varying levels of introgression or varying distances to ancestral populations, we performed two simulations on 1kb windows. For each statistic of interest we present the area under the curve (AUC) values. All simulations start with 10.000 loci under the neutral scenario ($f=0$) and 1.000 locus with subject to introgression. The Recombination rate is fixed at $r=0.01$.

Fraction of introgression: To test the impact of varying the fraction of introgression we simulated the fraction of introgression for the 'alternative model' from $f$=0.1 to $f$=1 and compared this to the neutral scenario where the fraction of introgression is zero ($f=0$). See below (supplementary table 6).

**Supplementary Table 6.** Fraction of introgression.

| Direction of gene-flow | Fraction of introgression | *D* | $f_d$ | *RNDmin* | $Bd_f$ |
|---|---|---|---|---|---|
| P3→P2 | 0.1 | 0.6252 | 0.7128 | 0.5579 | 0.7065 |
| P3→P2 | 0.2 | 0.6846 | 0.8426 | 0.6043 | 0.8330 |
| P3→P2 | 0.3 | 0.7163 | 0.9221 | 0.6479 | 0.9139 |
| P3→P2 | 0.4 | 0.7293 | 0.9541 | 0.7196 | 0.9478 |
| P3→P2 | 0.5 | 0.7380 | 0.9810 | 0.7588 | 0.9753 |
| P3→P2 | 0.6 | 0.7466 | 0.9922 | 0.8246 | 0.9890 |
| P3→P2 | 0.7 | 0.7585 | 0.9979 | 0.8510 | 0.9961 |
| P3→P2 | 0.8 | 0.7607 | 0.9988 | 0.9216 | 0.9980 |
| P3→P2 | 0.9 | 0.7748 | 1 | 0.9659 | 0.9996 |
| P3→P2 | 1 | 0.7871 | 1 | 1 | 0.9998 |

The effect of varying fractions of introgression on the model utility in the ROC analysis as indicated for values of AUC. The background history (coalescent times) is: $P12=1N_e$, $P123=2N_e$ and $P1234=3N_e$ generations ago. The calls to *ms* are:

*Neutral model: ms 32 1 -I 4 8 8 8 8 -ej 1 2 1 -ej 2 3 1 -ej 3 4 1 -r 10 1000*

*Alternative model: P3→P2: ms 32 1 -I 4 8 8 8 8 -ej 1 2 1 -ej 2 3 1 -ej 3 4 1 -es **0.1** 2 **Fraction** -ej **0.1** 5 3 -r 10 1000*

Distance of ancestral population:

Finally, by varying the distance to ancestral populations we tested the impact of a low amount of introgression on the various statistics.

**Supplementary Table 7.** Distance of ancestral population.

| Direction of gene-flow | Distance to ancestral population ( P12-P123-P1234) | *D* | $f_d$ | *RNDmin* | $Bd_f$ |
|---|---|---|---|---|---|
| P3→P2 | 0.3-1-3 | 0.6366 | 0.6860 | 0.5308 | 0.6782 |
| P3→P2 | 1-1-3 | 0.5884 | 0.6077 | 0.5107 | 0.5969 |
| P3→P2 | 0.5-2-3 | 0.6314 | 0.7292 | 0.5489 | 0.7272 |
| P3→P2 | 0.5-3-3 | 0.6454 | 0.7493 | 0.5498 | 0.7488 |
| P3→P2 | 1-2-3 | 0.6252 | 0.7128 | 0.5482 | 0.7065 |
| P3→P2 | 1-3-3 | 0.6465 | 0.7604 | 0.5690 | 0.7575 |
| P3→P2 | 2-2-3 | 0.6016 | 0.6406 | 0.5312 | 0.6220 |
| P3→P2 | 1.5-2-3 | 0.6065 | 0.6680 | 0.5526 | 0.6573 |

10.000 loci under the neutral scenario ($f$=0). Fraction of introgression for the 'alternative model' simulations is $f$=0.1 (1.000 locus). Recombination rate is r=0.01. Time of gene-flow is $0.1N_e$.

*Neutral model:*

*ms 32 1 -I 4 8 8 8 8 -ej **P12** 2 1 -ej **P123** 3 1 -ej **P1234** 4 1 -r 10 1000*

*Alternative model:*

*P3→P2: ms 32 1 -I 4 8 8 8 8 -ej **P12** 2 1 -ej **P123** 3 1 -ej **P1234** 4 1 -es 0.1 2 **0.9** -ej 0.1 5 3 -r 10 1000*

### 3 PopGenome Usage

```
# Install the PopGenome package from CRAN within R
install.packages("PopGenome")

# Install the PopGenome package from github
install.packages("dplyr")
install.packages("devtools")
library(devtools)
install_github("pievos101/PopGenome")

# Load the PopGenome package
library(PopGenome)
```

```
# Read the data (Fontaine et. al, 2015)
genome =
readVCF("AGC_refHC_bialSNP_AC2_2DPGQ.3L_V2.CHRcode2.DRYAD.vcf.gz",
10000,"4",1,45000000, include.unknown=TRUE)

# Define the populations
Aquad = c("SRS408143", "SRS408145", "SRS408151", "SRS408155",
"SRS408966", "SRS408969", "SRS408972", "SRS408973", "SRS408983",
"SRS420578")

Amela = c("SRS408142", "SRS408185", "SRS408994")

Ameru = c("SRS408186", "SRS408187", "SRS408967", "SRS408974",
"SRS408992", "SRS410266","SRS410284", "SRS410286", "SRS410290",
"SRS420577")

# Define the outgroup
Chris = c("CHRISTYI")

# Set the populations
genome = set.populations(genome, list(Aquad,Amela,Ameru),diploid=TRUE)

# Set the outgroup
genome  = set.outgroup(genome, Chris, diploid=TRUE)

# Transform the data into 50kb consecutive windows
slide = sliding.window.transform(genome,50000,50000,type=2)

# Perform the Bd-fraction, D and fd
slide = introgression.stats(slide)
# Perform the Bd-fraction with laplace smoothing
slide = introgression.stats(slide,  l.smooth=TRUE)

# Perform Pairwise Diversity
slide = diversity.stats.between(slide)
```

```
head(slide@nuc.diversity.between)
head(slide@BDF)

BDF = slide@BDF

head(slide@D)
head(slide@f)

D  = slide@D
f  = slide@f

# Get the genomic positions
genome.pos = sapply(slide@region.names, function(x){
split = strsplit(x," ")[[1]][c(1,3)]
val = mean(as.numeric(split))
return(val)
})

# Plot the results (Bd-fraction)
plot(genome.pos, BDF, pch=19, ylab="Bd-fraction", xlab="genomic
position", main="3La inversion")

# Perform the weighted jackknife for the inverted region
# First, split the data into 50kb windows within the inversion

slide =
sliding.window.transform(test,jump=50000,width=50000,start.pos=15000000,
end.pos=35000000, type=2)

# Perform the introgression statistics
slide  = introgression.stats(slide, l.smooth=TRUE)

# weighted jackknife
slide = weighted.jackknife(slide, per.region=FALSE)

# show results
slide@D.z
```

```
slide@D.pval
slide@BDF.z
slide@BDFD.pval

# Perform the drop-one jackknife for specific regions
# Split the data into the two regions 7.25kb-7.3kb and 21.85kb-21.90kb
split = splitting.data(test, positions=list(7250000:7300000,
21850000:21900000), type=2)

# Perform the statistics
split = introgression.stats(split,  l.smooth=TRUE)

split = weighted.jackknife(split, per.region=TRUE)
split@BDF
split@BDF.z
split@BDF.pval
```
